# Th17 T cells and immature dendritic cells are the preferential initial targets after rectal challenge with an SIV-based replication-defective dual-reporter vector

**DOI:** 10.1101/2021.04.27.441720

**Authors:** Danijela Maric, Wesley A. Grimm, Natalie Greco, Michael D. McRaven, Angela J. Fought, Ronald S. Veazey, Thomas J. Hope

## Abstract

Understanding the earliest events of HIV sexual transmission is critical to develop and optimize HIV prevention strategies. To gain insights into the earliest steps of HIV rectal transmission, including cellular targets, rhesus macaques were intra-rectally challenged with a single-round SIV-based dual reporter that expresses luciferase and iRFP670 upon productive transduction. The vector was pseudotyped with the HIV-1 envelope JRFL. Regions of tissue containing *foci* of luminescent, transduced cells were identified macroscopically using an *in vivo* imaging system, and individual transduced cells expressing fluorescent protein were identified and phenotyped microscopically. This system revealed that anal and rectal tissues are both susceptible to transduction 48 hours after the rectal challenge. Detailed phenotypic analysis revealed that on average, 62% of transduced cells are CCR6^+^ T cells—the vast majority of which express RORγT, a Th17 lineage-specific transcription factor. The second most common target cells were immature dendritic cells at 20%. These two cell types were transduced at the rates that are four to five times higher than their relative abundances indicate. Our work demonstrates that Th17 T and immature dendritic cells are preferential initial targets of HIV/SIV rectal transmission.

**IMPORTANCE:** Men and women who participate in unprotected receptive anal intercourse are at high risk for acquiring HIV. While *in vitro* data have developed a framework for understanding HIV cell tropism, the initial target cells in the rectal mucosa have not been identified. In this study, we identify these early host cells by using an innovative rhesus macaque rectal challenge model and methodology, which we previously developed. Thus, by shedding light on these early HIV/SIV transmission events, this study provides a specific cellular target for future prevention strategies.

## INTRODUCTION

Receptive anal intercourse (RAI) accounts for more than half of new HIV-1 infections in the United States [1]. For a successful transmission event to occur, HIV has to overcome physical and biological barriers—including mucus, epithelia and host restriction factors. While the initial events in vaginal transmission were explored by us [2, 3] and others [4-6] in several studies, little is known about HIV infection via intrarectal route. The goal of this study was to identify the initial cell targets of HIV infection during rectal transmission.

Epidemiological and comparative studies in macaque rectal challenge models suggest that HIV/SIV is more efficiently transmitted via RAI than via other mucosal routes [7, 8]. In RAI, incoming virions encounter anal and rectal tissues, which are structurally distinct [9]. Anal tissue is composed of a stratified multilayered epithelium, which is similar to vaginal and ectocervical tissue. In contrast, rectal tissue, like endocervical, is lined by a simple columnar epithelium (i.e., a single cell layer). It is generally believed that the stratified squamous epithelium provides a more robust protection against invading pathogens relative to the single cell layer of the columnar epithelium, which is expected to be more readily penetrated by external solutes and microorganisms. In addition, underlying the simple epithelium are a large number of CD4^+^ HIV-target cells, including: T cells, macrophages and dendritic cells (DCs) [10], further increasing the susceptibility of the rectum to infection by HIV/SIV relative to the oral mucosa and female reproductive tract (FRT). There is an ongoing debate about the initial targets of infection after rectal transmission. Some reports suggest that T cells are the earliest cell targets of HIV/SIV infection, while others support the initial infection of macrophages [11, 12]. A trojan horse model where DCs pick up the virus at the mucosal site of exposure and ferry it to draining lymph nodes has also been proposed [13]. A definitive answer to this debate is still pending. Clarifying the earliest targets of HIV infections is critical to understand HIV rectal transmission and for the development and optimization of HIV prevention strategies.

We previously developed a dual-reporter system to study early HIV transmission events in the FRT [2]. Using this reporter system and the rhesus macaque (*Macaca mulatta*; RM) vaginal challenge model, we were able to survey the entire FRT for sites of infection (i.e., luciferase^+^ cells) and to phenotype the initial target of infection. By 48 hours post challenge, we detected infected cells in both lower and upper FRT—including vagina, ecto- and endocervix, and ovaries. Importantly, in all FRT tissues, the vast majority of infected cells were CD4^+^ T cells [2]. Subsequently, using replication competent SIV_mac_239, we identified these early targets to be Th17 T cells [3].

To study HIV transmission after rectal exposure and determine whether vaginal and rectal transmission share similar initial events, we developed a replication-defective version of the dual-reporter system used in our FRT studies. This dual-reported SIV was used to challenge rhesus macaques. All transduced cells identified in the exposed rectal mucosa represent infection by the challenge inoculum. Thus, these cells are equivalent to the first cells infected in the tissue after sexual exposure. Our data reveal that Th17 and immature dendritic cells (iDCs) are the earliest cells to be infected/transduced after rectal exposure. Importantly, transduced cells were detected throughout the rectum and anus—revealing that both the columnar epithelium of the rectum and the stratified squamous epithelium of the anus are permissive to HIV/SIV infection.

## RESULTS

### Dual-reporter lentiviral vector: design, validation and production

Our laboratory developed an SIV-based dual-reporter replication-defective vector encoding luciferase and mCherry [14] proteins to identify the initial viral targets in the vaginal-challenge RM model [2]. The luciferase reporter allows for low-resolution screening to identify small *foci* of transduced, luciferase expressing cells in large pieces of tissue. The luciferase-positive *foci* can then be dissected, cryosectioned and studied by, for example, fluorescent microscopy and PCR.

For the current study, we replaced mCherry with the near-infrared fluorescent protein 670 (iRFP670). The iRFP670 fluorophore is better for imaging tissues with high autofluorescence (e.g., mucosal tissues in general and rectal tissue especially), because of its brightness and narrow emission spectrum [15]. In addition, there is less autofluorescence at longer (near-infrared) wavelengths [16]. Expression of this modified bicistronic construct was driven by the CMV immediate-early promoter (Fig. 1A). While the luciferase gene [17] is expressed via translation of the 5’-most initiation codon, iRFP670 was translated via an internal ribosome entry site (IRES) [2]. Lastly, SIV long terminal repeats (LTR) flank the dual-reporter expression cassette. We refer to this vector system as “**LI670”**—based on the sequential **<u>luciferase</u>, <u>I</u>**RES and iRFP**<u>670</u>** cassettes.

**FIG 1.**
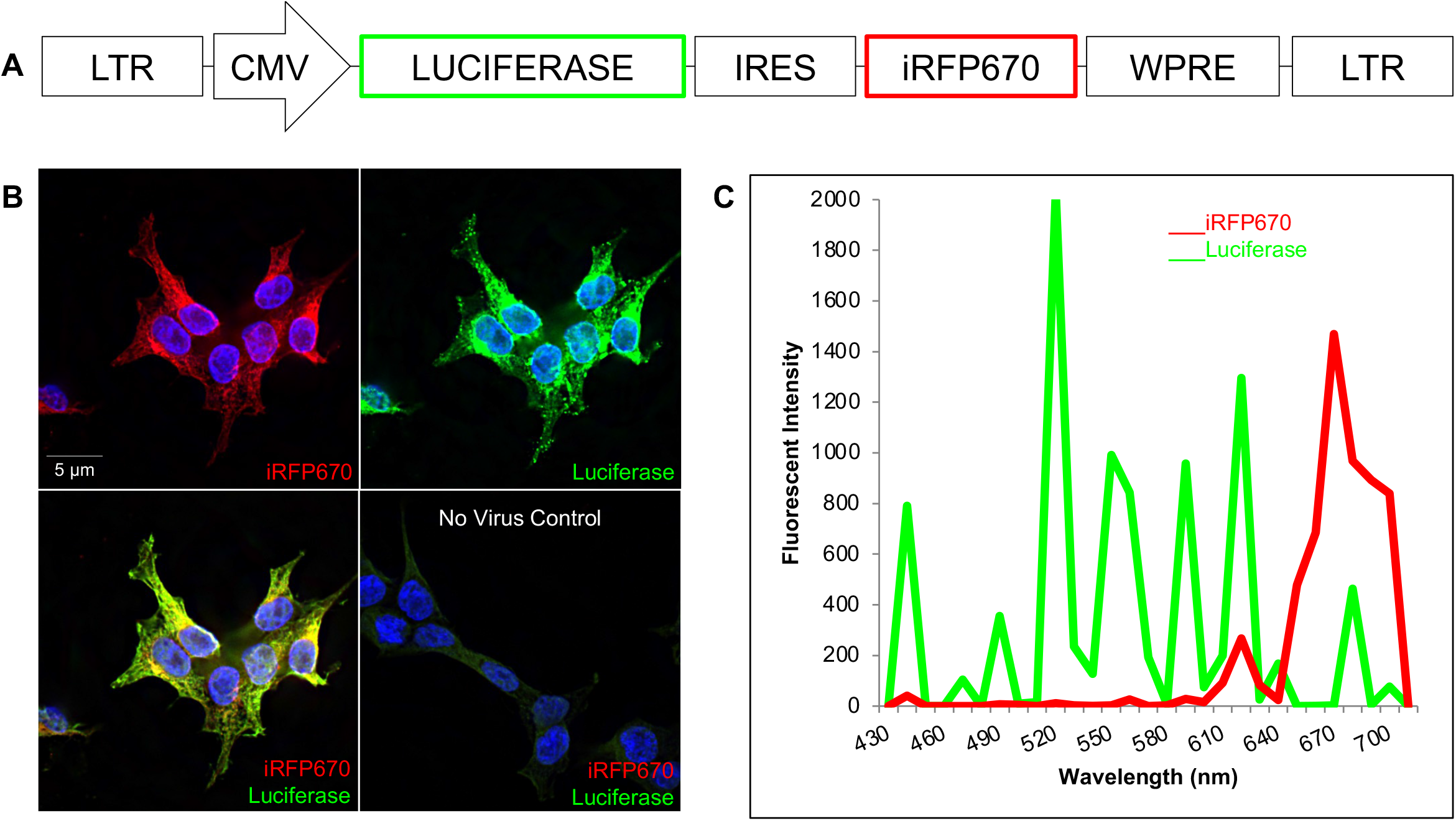
Design and in vitro validation of dual-reporter lentiviral vector. (A) LI670 lentiviral reporter vector contains luciferase and iRFP670 genes driven by CMV and IRES promoters, respectively. (B) 293T cells transduced with VSVg-LI670 virus express iRFP670 (red) and are immunopositive for luciferase (green); nuclei labeled by Hoechst stain (blue). Untransduced, negative-control 293T cells. (C) Spectral emission profile of a transduced 293T cell.

To validate the LI670 construct in vitro, 293T cells were transduced with VSVg-pseudotyped particles (VSVg-LI670) and, after 48 hours, fixed, immunolabeled for luciferase and analyzed by fluorescence microscopy. As expected, all iRFP670^+^ cells were also immunopositive for luciferase, and had clearly defined Hoechst-stained nuclei (Fig. 1B). Un-transduced 293T cells had no luciferase or iRFP670 signal above background. To validate that the detected fluorescent signals were due to iRFP670 and Alexa Fluor 488-labeled luciferase antibody, we performed spectral confocal imaging; this allows the unique fluorescent emission signature of each fluorophore to be identified, and distinguished from autofluorescent signals. In fact, the maximum emissions of the transduced 293T cells precisely matched those anticipated— very narrow emission spectra with maxima at 670 nm and 520 nm for iRFP670 and Alexa Fluor 488, respectively (Fig. 1C).

To produce viral particles for the rectal challenge experiments, 293T cells were co-transfected with the LI670 vector and a SIV3 packaging construct. This provided the viral proteins in trans to package the LI670 genomic RNA, as we previously described [2]. The resulting LI670 particles were pseudotyped with the HIV JRFL envelope [18] (JRFL-LI670), which directs the particles to fuse with CD4^+^ and CCR5^+^ cells. The LI670 replication-defective dual-reporter particles do not encode any viral proteins. Thus, they transduce target cells, but do *not* produce viral particles. As a result, the only cells transduced after viral inoculation *in vivo* are the ones initially targeted by the challenge inoculum. These cells are equivalent to the earliest cell targets after HIV rectal infection.

### Both anal and rectal tissue are susceptible to transduction by HIV/SIV after rectal challenge

To investigate the earliest events after rectal transmission, four female RMs were inoculated intrarectally with 5 mls of concentrated JRFL-LI670 virus (TCID_50_ range: 10^4^-10^6^). Prior to viral challenge, we collected rectal biopsies to mimic the effect of microtrauma that would result from unprotected RAI, and to provide a positive reporter signal as a result of focal transduction. Animals were sacrificed 48 hours post challenge.

Anal and rectal tissue are structurally and functionally distinct. Our IVIS-luciferase system allowed us to rapidly assess the distribution of transduced cells over a large surface area and determine whether rectal and anal tissues also differ in their susceptibility to HIV infection.

Immediately following sacrifice, the entire anus and part of the adjoining rectum were removed in one piece, washed and examined by IVIS to assess background luminescence. Minimal to no background luminescence was noted in each of four animals (Figs. 2A, S1A). Tissue at each visible biopsy site was excised to serve as positive control. The remaining tissue was cut into six pieces. All the pieces were incubated in luciferin and re-examined by IVIS to detect luciferase expression. The number, size and distribution of luciferase-positive *foci* varied between animals (Figs. 2B, S1B)—in a way reminiscent of our vaginal challenge studies [2]. One of the four animals showed substantial luciferase signal throughout the anorectal tissue (FN94; Fig. 2B). In other two animals the luciferase signal was concentrated in the anal region (GG70 and HP63; Fig. S1B). In contrast, the fourth animal had minimal to no luciferase signal (DK09; Fig. 2B). The tissue excited from the site of the biopsies exhibited robust luciferase signal in all animals except in HP63, where the signal was minimal (Figs. 2B, S1B). In three of the four animals, luciferase signal was strong in anal tissue. This was unexpected given that the stratified epithelium of the anal tissue is thought to constitute a robust barrier against pathogens and it suggests that the anal and the rectal tissues are both susceptible to HIV infection.

**FIG 2.**
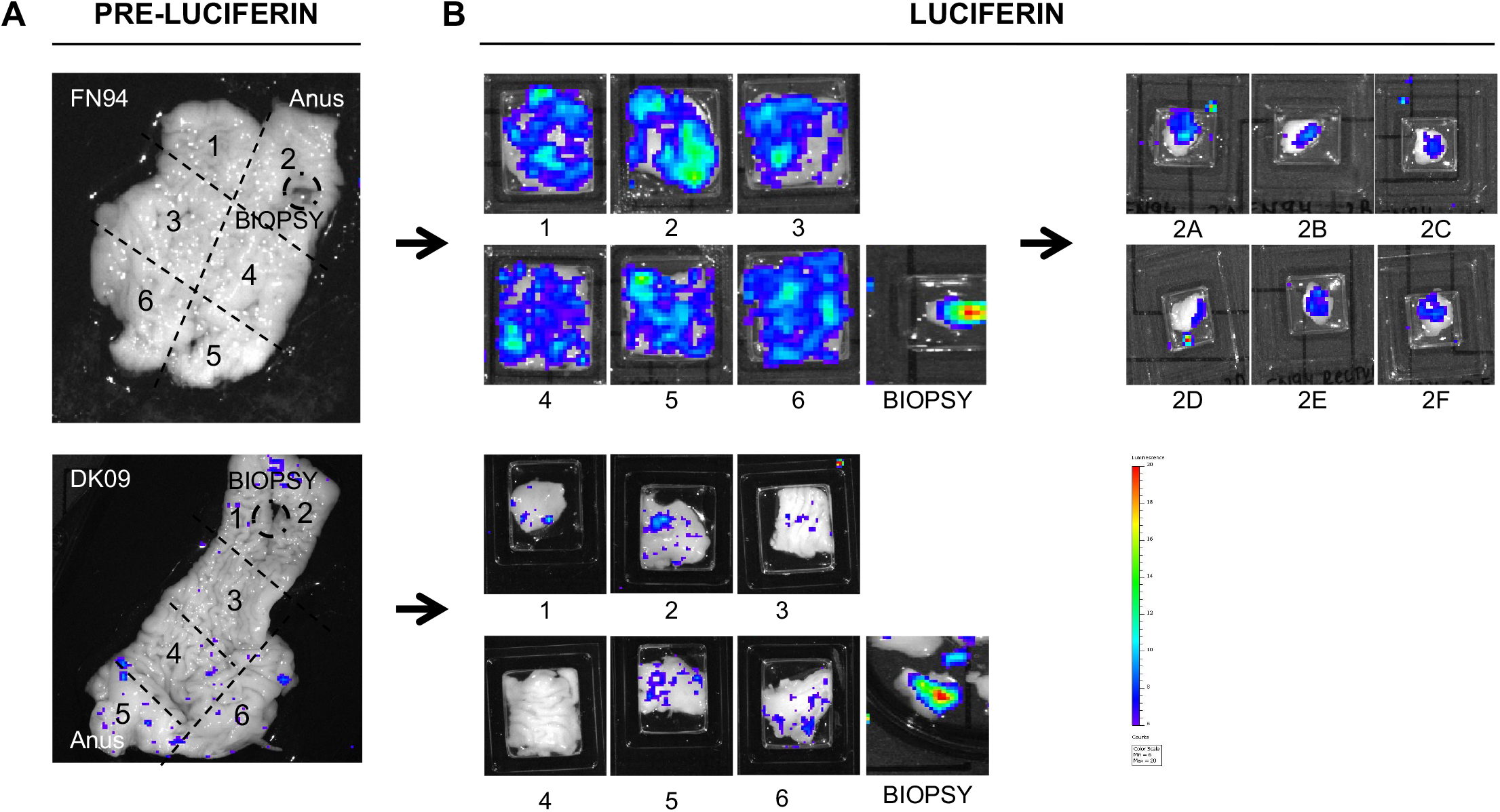
Luciferase reporter expression in anal and rectal tissue of vector-inoculated RMs. Animals were inoculated with JRFL-pseudotyped dual-reporter vector, sacrificed 48 hours later and the first 6 to 7 cm of distal colon removed in one piece. (A) Extirpated anorectal tissue from two animals (FN94 and DK09) imaged by IVIS to assess background luminescence; biopsy sites and anal pole are indicated. (B) Tissue at the biopsy sites was excised and remaining tissue cut into six large pieces as indicated by dotted lines in A; all pieces were soaked in d-luciferin and imaged by IVIS to visualize luciferase expression. For animal FN94, the tissue piece with the highest expression was cut into six smaller pieces that were then re-imaged by IVIS.

To validate that the luciferase-positive *foci* detected by IVIS represent transduced cells, regions of tissue with the strongest luciferase signal were flash-frozen, cryosectioned, immunolabeled, and imaged by fluorescent microscopy. E-cadherin immunostaining in the green channel (Alexa Fluor 488) revealed the distinct epithelial gross morphologies: the stratified epithelium of anus and the simple columnar epithelium of rectum (Fig. 3A). Imaging in the far-red channel to detect iRFP670 revealed bright red *puncta* that were most commonly found in the *lamina propria* of both tissues (Fig. 3A). Analysis at higher magnification showed that these red *foci* were tightly associated with Hoechst-stained nuclei—essentially enveloping the nuclei (3 insets in Fig. 3A and 3B). Sections were then stained with an anti-luciferase antibody in Alexa Fluor 488 to confirm the colocalization of the iRFP670^+^ signal and luciferase. We examined 53 iRFP670^+^ cells across all four animals, and they all expressed luciferase (Fig. 3B). We next validated that the observed fluorescence was due to the iRFP670 and Alexa Fluor 488 tagged luciferase, using spectral imaging and a scanning laser confocal microscope (Fig. 3C). As expected, the emission profiles of the putatively transduced cells matched the known profiles and emission maximums maxima for iRFP670 and Alexa Fluor 488.

**FIG 3.**
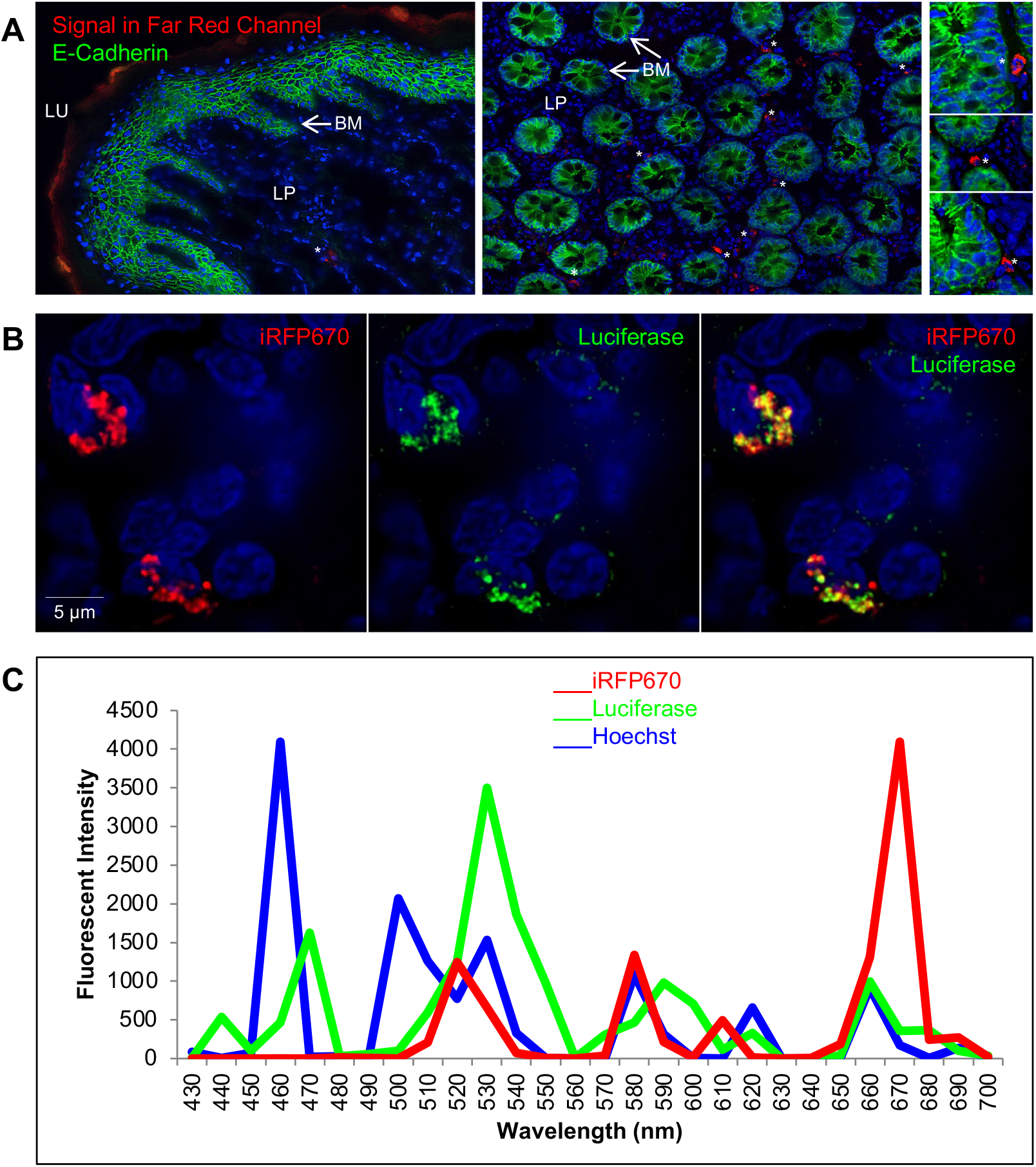
Identification of JRFL-LI670 transduced cells in the anorectal tissue. (A) E-cadherin immunostaining (green) of a longitudinal cryosection of the anal tissue (left), and of a transverse section through the upper part of the intestinal glands of the rectal tissue (right)—including three high magnification inserts. iRFP670^+^ cells (red puncta, asterisks); Hoechst nuclear stain (blue); BM, basement membrane; LP, lamina propria; LU, lumen. (B) Single-cell colocalization of iRFP670 fluorescent reporter protein (red) and luciferase immunostaining (green); Hoechst stain (blue). (C) Spectral profiles and emission peaks of iRFP670 (red, 670 nm), Zenon Alexa Fluor 488-labeled luciferase antibody (green, 530 nm) and Hoechst stain (blue, 460 nm).

The ability of the IVIS/luciferase assay to detect transduced cells is dependent on the size and depth of the luciferase-positive *foci* [2]. To determine if tissue areas that appear luciferase-negative by IVIS actually contain transduced cells, we carried out nested PCR—a more sensitive methodology that we previously used to detect rare transduction events in FRT tissues [2, 19, 20]. DNA was isolated from cryosections of three different types of tissue: 1) two Optimal Cutting Temperature (OCT) media blocks of rectal tissue from animal DK09 that were luciferase-negative by IVIS, 2) tissue from a biopsy site in animal DK09 (positive control) and 3) anorectal tissue from an unchallenged animal from unrelated study (negative control). As expected, all twelve lanes of positive-control DNA exhibited a robust 244-bp band—a fragment of the luciferase gene (Fig. S2), and negative-control DNA had no detectable signal. Interestingly, the luciferase-target band was present in 6 of 24 lanes containing DNA from tissue that was luciferase-negative by IVIS (Fig. S2). This suggests that some of the areas of tissue that appear negative in the IVIS-luciferase images contain transduced cells.

### Th17 CD4^+^ T cells are the preferential first target of HIV infection after rectal challenge

Th17 cells are preferentially infected by HIV in vitro [21, 22] and preferentially depleted in HIV-infected patients and SIV infected macaques [23, 24]. Moreover, we have previously shown that Th17 T cells are the preferred early target of SIVmac239 in FRT of RM and that dual expression of CD3^+^ and CCR6^+^ is a strong predictor of Th17 cell phenotype [3]. Thus, we hypothesized that Th17 may be among the earliest target of HIV/SIV infection after rectal transmission as well. To determine if this was the case, we immunostained the cryosections of luciferase-positive anorectal tissue from the four macaques rectally challenged with JRFL-LI670 with antibodies against one or more of three cell surface receptors: CD4 (HIV-1 receptor), CD3 (T-cell-specific marker) and CCR6, a chemokine receptor that is expressed on iDCs and Th17 cells [25, 26].

We found that all iRFP670^+^/luciferase^+^ cells were CD4^+^, as expected, and that the majority (69% ± 11%) were CD3^+^ (Fig. 4A and 4B; Table S1). This was consistent with our previous vaginal challenge studies [2]. Double immunolabeling for CD3 and CCR6 (Fig. 4C; Table 1) revealed that the majority (62% ± 8%) of transduced cells were CD3^+^/CCR6^+^ (ie, likely Th17 cells); the second highest percentage (20% ± 8%) was CD3^−^/CCR6^+^ cells (ie, likely iDCs). These results suggest that the predominant target of JRFL-pseudotyped virion in anorectal tissue are Th17 CD4^+^ T cells and, to a lesser extent, iDCs. The two other target cell phenotypes examined were transduced at lower frequency: CD3^+^/CCR6^−^ (non-Th17 T cells; 4% ± 5%) and CD3^−^ /CCR6^−^ (other cells; 14% ± 14%). Since at least one of the four averaged proportions was different (*p* < .01), two pairwise comparisons were performed: 1) CD3^+^/CCR6^+^ was greater than CD3^+^/CCR6^−^, CD3^−^/CCR6^+^ and CD3^−^/CCR6^−^; and 2) CD3^+^/CCR6^−^ was lower than CD3^−^/CCR6^+^ (all *p* values less than the Bonferroni corrected alpha of 0.008).

**FIG 4.**
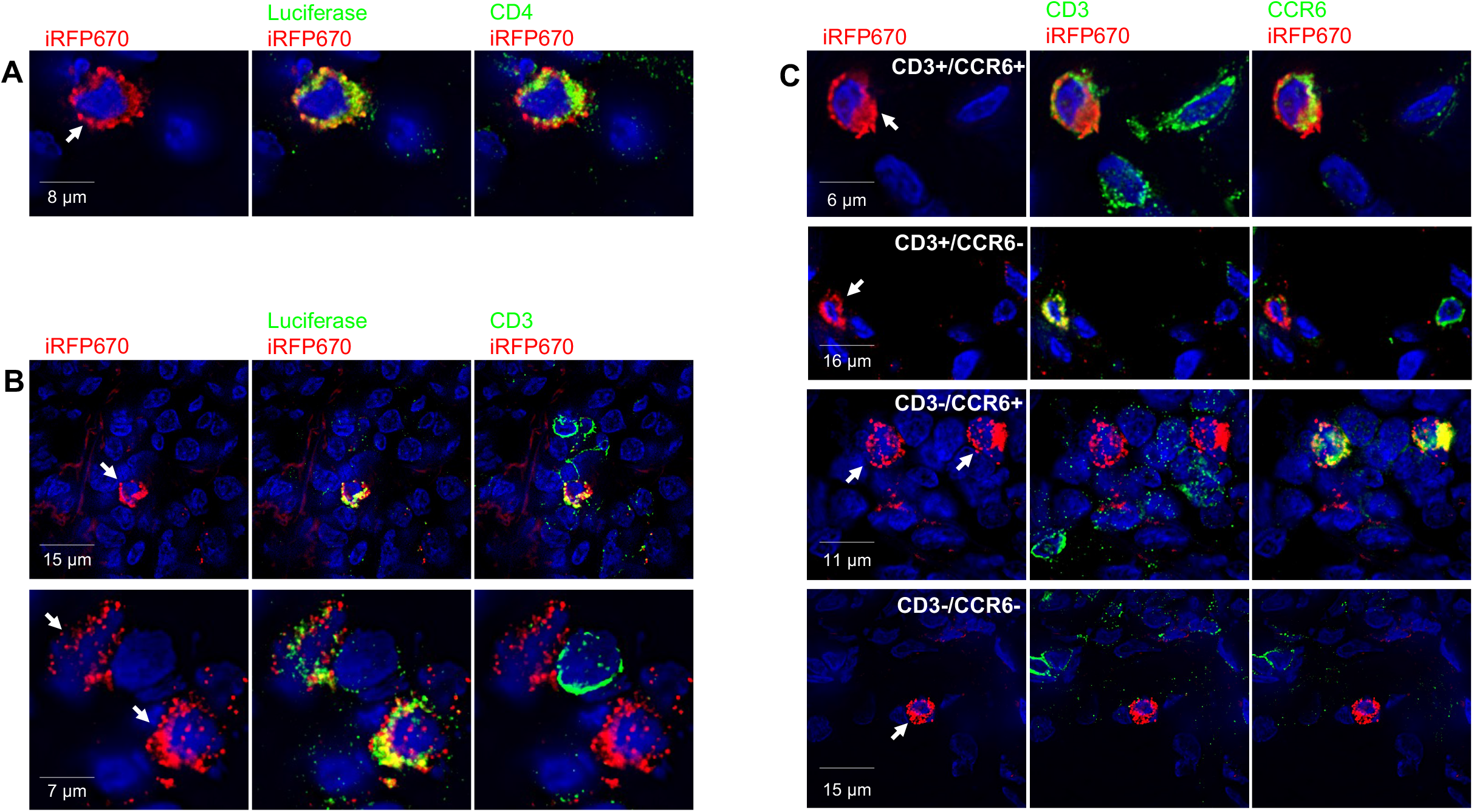
Four distinct phenotypes of early transduced cells in anorectal tissue. Cryosections were immunolabeled (green) for one or more cell surface markers (CD4, CD3 and CCR6) and, in some instances, luciferase. iRFP670, fluorescent reporter protein (red); Hoechst stain (blue). Each row shows three images of the same field. (A) CD4^+^ transduced cell (arrow). (B) CD3^+^ transduced cell (arrow, top) and CD3^−^ transduced cells (arrows, bottom). (C) Dual immunostaining for CD3 and CCR6 reveals four distinct phenotypes of transduced cells (arrows).

**Table 1.** CD3 and CCR6 phenotypic analysis of transduced* cells after anorectal inoculation with JRFL-pseudotyped virions.

| Animals | Cell Count (%) |  |  |  |  |
| --- | --- | --- | --- | --- | --- |
|  | CD3<sup>+</sup> |  | CD3<sup>-</sup> |  | TOTAL |
|  | CCR6<sup>+</sup> | CCR6<sup>-</sup> | CCR6<sup>+</sup> | CCR6<sup>-</sup> |  |
| GG70 | 6 (67%) | 0 (0%) | 2 (22%) | 1 (11%) | 9 |
| HP63 | 3 (50%) | 0 (0%) | 1 (17%) | 2 (33%) | 6 |
| FN94 | 17 (65%) | 1 (4%) | 8 (31%) | 0 (0%) | 26 |
| DK09 | 6 (67%) | 1 (11%) | 1 (11%) | 1 (11%) | 9 |
| **Total (% average ± SD)** | **32 (62% ± 8%)** | **2 (4% ± 5%)** | **12 (20% ± 8%)** | **4 (14% ± 14%)** | **50** |
| * iRFP670<sup>+</sup> |  |  |  |  |  |

While dual expression of CD3 and CCR6 is commonly used to identify Th17 cells, a more conclusive marker is RORγT, a master regulator transcription factor that controls IL-17 production in T cells [25]. Therefore, to validate that the CD3^+^/CCR6^+^ T cells noted in anorectal mucosa are, in fact, Th17 cells, we tested for expression of RORγT. Since no antibodies are available to detect the RORγT, we utilized fluorescent RNA probes and in situ hybridization (RNA-FISH), as we have done previously in the FRT [3]. RORγT FISH was optimized (Fig. 5A) and validated by spectral imaging (Fig. 5B). To visualize RORγT mRNA, CD3 and CCR6 in the same cells, cryosections were processed for RNA-FISH (RORγT) and immunofluorescence (IF; CD3 and CCR6) (Fig. 5C). Thus, in this analysis, we evaluated all cells, blind to transduction status. We found that the vast majority (roughly 90%) of CD3^+^/CCR6^+^ cells in the rectal tissue expressed RORγT. These data validate that dual expression of CD3 and CCR6 is a strong indicator of the Th17 cell phenotype in RM anorectal tissue and has provided further evidence that Th17 T cells represent the majority of the early cellular targets of HIV/SIV following the rectal challenge.

**FIG 5.**
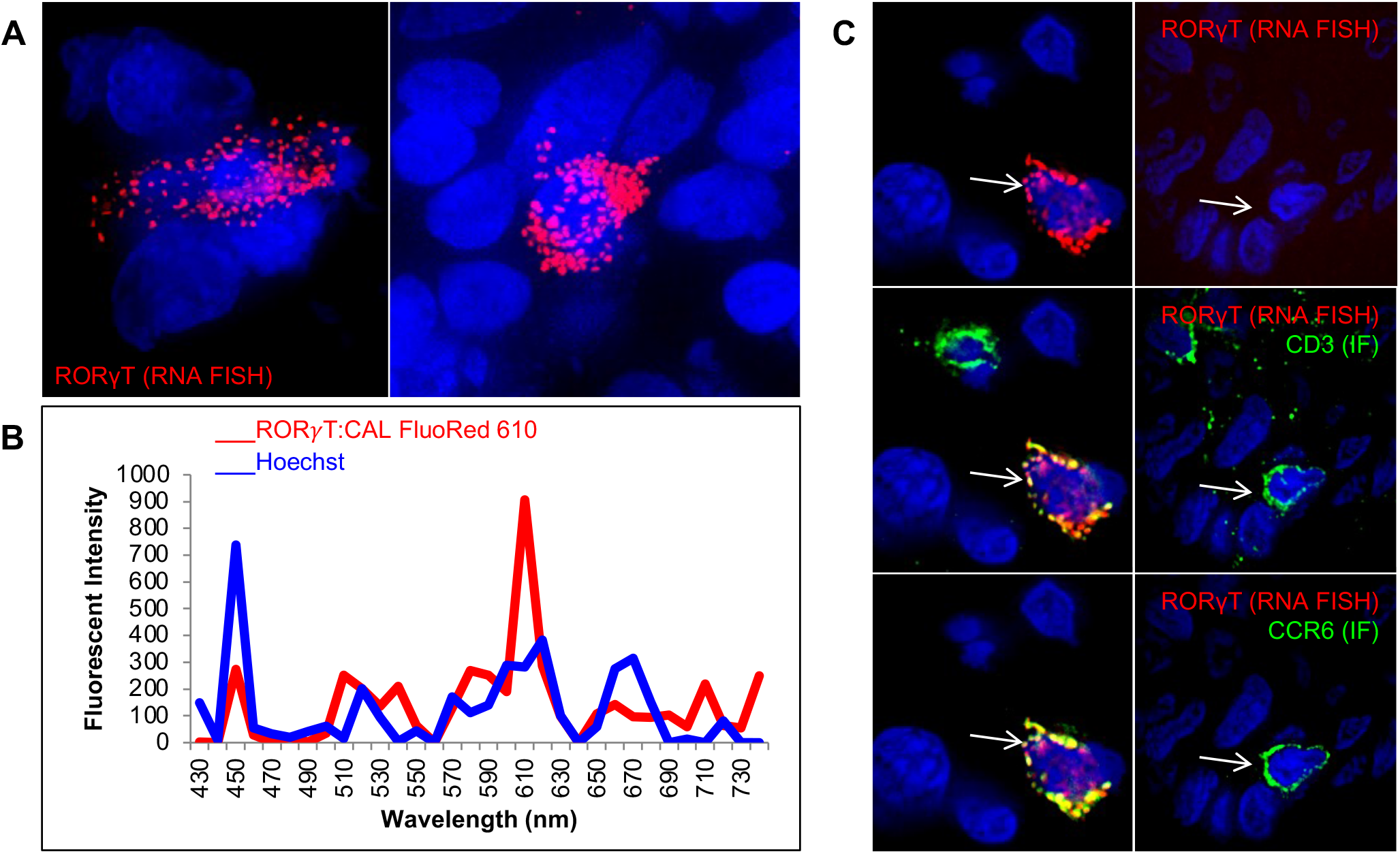
The majority of CCR6^+^ CD3^+^ T cells express RORγT in the anorectal tissue. RM anorectal tissue was processed for RORγT FISH (red) and/or IF for CD3 and CCR6 (green). Hoechst nuclear stain (blue). (A) RORγT FISH. (B) Spectral imaging of one of the RORγT^+^ cells in A. CAL Fluo Red 610-labeled RORγT (red, 610 nm) and Hoechst stain (blue, 460 nm). (C) Sequential IF and FISH; CD3^+^/CCR6^+^ cells that are RORγT^+^ (left panels, arrows) or RORγT^−^ (right panels, arrows).

In addition to Th17 T cells, the CCR6 chemokine is also expressed on iDCs [26]. Our phenotypic analysis revealed that the second most predominant population of transduced cells are CD3-/CCR6+ cells accounting for 20% ± 8%. To find out if CD3-/CCR6+ cells at RM anorectal tissues are dendritic cells we assessed the expression of DC-SIGN molecule on these cells by adding the CD209 antibody to the staining protocol. The triple labeling with CD3, CCR6 and CD209 antibodies reveled that all of the CD3-/CCR6+ cells expressed CD209 molecule and were hence classified as iDCs (Fig. S3).

### The preferred HIV-target cells occur at low frequencies in anorectal tissue

Th17 T cells and, to a lesser extent, iDCs were the cell types that were most commonly targeted by our JRFL-pseudotyped LI670 vector (Table 1). To test whether this is simply a reflection of their relative abundance in anorectal mucosa, we analyzed the frequencies of the same four CD4^+^ cell phenotypes regardless of transduction status. To do this, cryosections from all four RMs were triple immunolabeled for CD4, CD3 and CCR6 (Figs. 6, S4). Triple immunolabeling revealed CD4^+^ cells scattered throughout the anorectal tissue (Fig. S4). Higher magnification imaging revealed that the four cell phenotypes (based on marker analysis) had distinguishing morphological characteristics (Fig. 6A). Th17 T cells and non-Th17 T cells were small and round, iDC cells were small with multiple cell projections, and the cells in the “other” category were larger and myeloid-like. We then quantified the relative frequencies of each of the four phenotypes: non Th17 T cells (71% ± 9%), other cells (15% ± 4%), Th17 T cells (12% ± 5%), and iDCs (2% ± 1%) (Fig. 6B, Table S2). Thus, in our model system, Th17 cells and iDCs, the predominant early targets, were the least abundant of the four CD4^+^ cell types. In fact, comparing these relative mean percentages with those of the iRFP670^+^, transduced cells (Table 1) revealed that Th17 T cells and iDCs were transduced at rates four to five times greater than would be expected if targeting were solely a function of relative abundance (Fig. 6B). In summary, our data demonstrate that Th17 T cells and iDCs are the preferential initial targets for SIV/HIV in macaque rectal challenge model.

**FIG 6.**
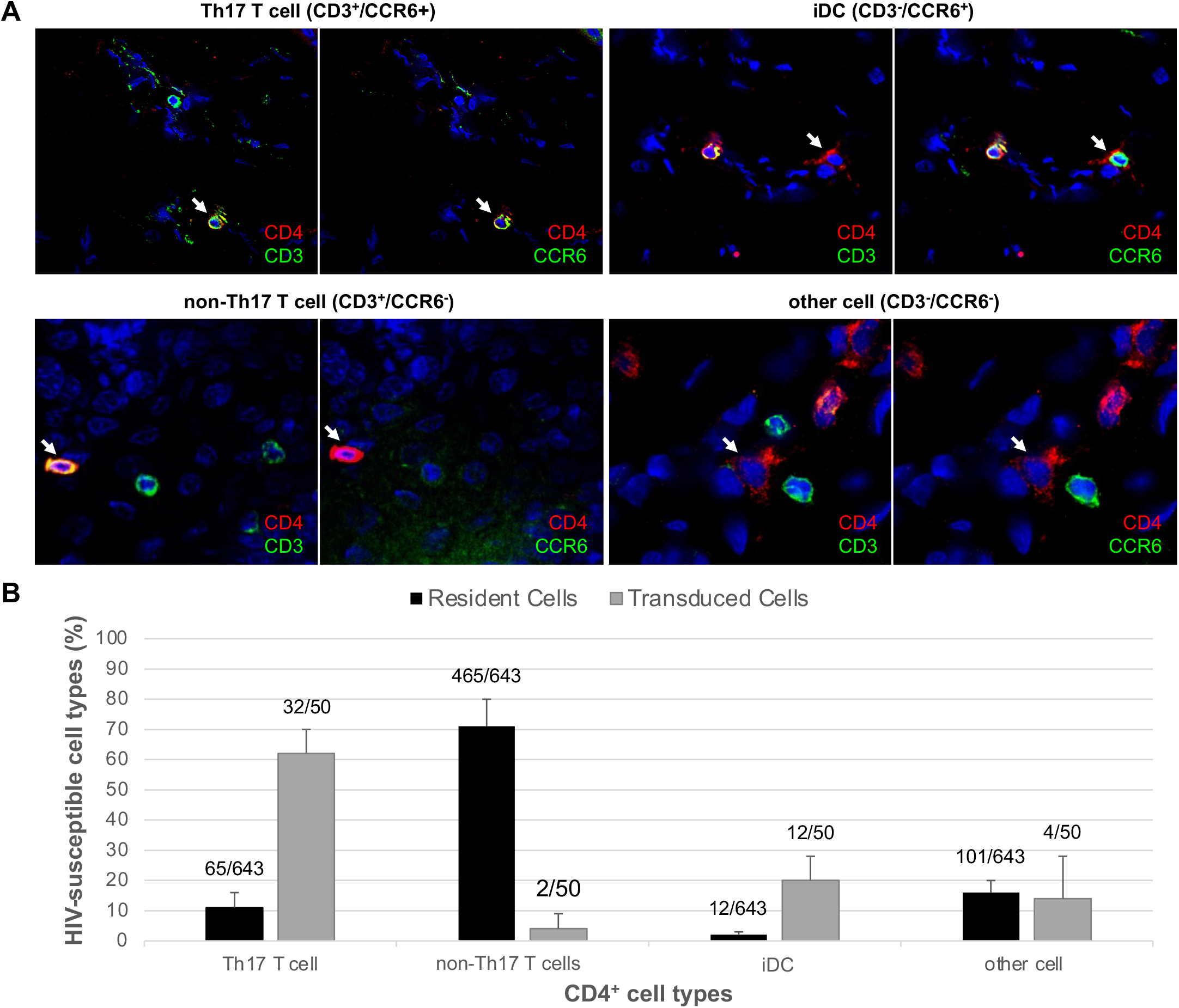
Th17 T cells and iDCs are transduced at five times greater frequency than their relative abundance would predict. Cryosections of anorectal tissues were triple immunolabeled for CD4, CD3 and CCR6, revealing four different CD4^+^, HIV-susceptible cell types (arrows in A). (A) Representative fluorescent images from four cryosections. CD4 (red), CD3 (green; far-red channel) and CCR6 (green; green channel); Hoechst stain (blue). (B) The four cell phenotypes displayed as a percentage of the 643 total CD4^+^ cells identified (black bars); data are from all four animals. Juxtaposed are the data from Table 1—the four cell phenotypes displayed as a percentage of the 50 total iRFP670+ (transduced) cells identified (gray bars). Error bars represent standard deviation.

## DISCUSSION

The overarching goal of this study was to gain a better understanding of the earliest stages of HIV infection during rectal transmission—specifically, to define the locations and phenotypes of the earliest target cells. To accomplish this, we used a modified version of our dual-reporter vector (JRFL-LI670), which we previously validated in a vaginal challenge RM model [2, 3]. Using the first reporter, luciferase, as a beacon, we were able to identify discrete *foci* of transduced cells within large areas of anorectal tissue by IVIS. Using the second reporter iRFP670, we were able to identify and characterize the transduced cells by fluorescent microscopy.

We validated the modified viral vector *in vitro* and *in vivo*. Nested PCR was used to detect the dual-reporter genome in anorectal tissues of infected macaques. Immunofluorescent microscopy (IF) was used to validate the expression of both reporter proteins (immunostaining for luciferase expression and direct visualization of iRFP670 fluorescent protein) along with spectral analysis. Lastly, we confirmed that the transduction was exclusive to CD4 positive cells.

With this system we were able to phenotype the preferred target cells of the challenge inoculum. As expected, we found that CD4^+^ T cells constituted the majority of the transduced cells. However, we also determined that most of these transduced CD4^+^ T cells presented a Th17 phenotype. This is despite the fact that Th17 constitute a relatively low proportion of CD4^+^ T cells in the anorectal tissue.

Interestingly, the second most abundant target was not another T cell subset, rather it was iDCs (ie, CD3^−^/CCR6^+^), which also occur at a very low frequency in the anorectal tissue. Early infection of iDCs support the hypothesis that these cells may act as trojan horses and facilitate viral transport to lymphoid sites increasing viral spread after transmission.

These data bear striking similarities to our vaginal transmission studies, using either LICh (dual-reporter vector with mCherry) [2] or wild-type SIVmac239 virus [3]. In both studies, CD4^+^ T cells were preferentially targeted in the 48 hours after challenge. Moreover, the wild-type SIV study, like the current study, revealed a four- to five-fold enrichment in infection of Th17 cells, which also constituted the vast majority of infected cells. This preference for Th17 cells in both vaginal and rectal mucosa may be explained by co-receptor usage. Thus, it may be important to determine whether Th17 in the rectal mucosa express higher levels of CCR5 than other T cell subsets. Moreover, future studies will need to investigate whether our data are recapitulated using a CXCR4-tropic envelope. Besides co-receptor expression, Th17 cells may express other surface molecules that may facilitate viral entry and/or transduction. In contrast, preferential transduction of iDCs may be due to their remarkable ability to capture viral particles through DC-SIGN or other lectin receptors, expressed at high levels on iDCs [13]. However, since these receptors and co-receptors are expressed at various levels also by other cell subsets, preferential transduction may be more readily explained by the state of activation of the cells. A higher metabolic state may facilitate nuclear transport, integration and transcription of our vector as well as of HIV. Th17s are involved in maintaining epithelial homeostasis by constitutively monitoring the levels of local commensal antigens presented by iDCs [27, 28]. This ongoing activity and regular function of these two cell types are consistent with them normally being in a more metabolically active state than other resident T cell subsets and likely provide an intracellular environment that efficiently supports the early phase of the SIV/HIV replication cycle [29]. This model is supported by data demonstrating that increased metabolic activity in target cells leads to increased reverse transcription, integration and virion production and that HIV does not efficiently replicate in resting CD4^+^ T cells [30, 31]. Alternatively, Th17 cells and iDCs may have lower expression of restriction factors such as Mxß, which can inhibit the early phase of the HIV life cycle, especially in a type-1 interferon-influenced environment [32, 33]. Therefore, susceptibility mediated by cellular metabolism or innate responses is potentially a more important determining factor than tropism in the sexual transmission of HIV.

This study focused on the anus and distal-most 6 cm of the rectal compartment and, therefore, did not examine transmission events that may have taken place in the descending or transverse colon. Fluids introduced into the rectum can rapidly spread beyond the distal rectum [34, 35] including as far as the distal (sigmoid) colon [35, 36]. This has been confirmed in the RM model and shown to be volume dependent, utilizing including methylene blue dye and MRI detectable viral surrogates [37] and the spectrum of target cells in mucosa can vary in different parts of the large bowel [38]. In addition, the atraumatic inoculation utilized here is distinct from receptive anal intercourse—both in terms of the resulting tissue trauma and the presence of semen, which could influence the location and phenotype of transduced cells. For example, receptive anal intercourse results in changes in the microbiome and increases in Th17 cell signatures [39]. Future studies are needed to address each of these concerns and to determine if they affect the localization and phenotypic profile of the initial cell targets.

Of note, our method did not efficiently detect isolate transduced cells in the anorectal tissue of the macaques. This is likely because photons used for bioluminescence (luciferase) detection are limited in the ability to penetrate the tissue. In contrast, nested-PCR allowed the detection of luciferase in a small percentage of tissues negative at IVIS. Since, the IVIS-negative, PCR-positive tissue represented only a fraction of the total PCR-positive tissue, our results would likely not change if these areas were included in our analysis. However, alternative imaging approaches such as radioactivity-based imagining by positronic emission tomography (PET) may be more sensitive for this type of studies.

Another downside of our system is the use of a replication incompetent virus that does not allow to study virus-host cell response to infection nor changes in the phenotype of infected cells over time. Thus, future studies will include an inoculum containing a mixture of our LI670 vector and wild-type SIV virus. We have already shown that this approach facilitates the identification of cells infected by the wild-type virus [3].

Future work will also have to address how risk factors, such as sexually transmitted infections and inflammatory microbiomes, that are known to impact the susceptibility of mucosal HIV transmission change the phenotype of the earliest target cells. Understanding the mechanisms of HIV transmission in receptive anal intercourse constitutes an important step to develop better prevention strategies. An increased understanding of the similarities and differences of the initial targets of infection in the vaginal and rectal compartments may advance approaches that increase protection in both compartments. The rhesus macaque model and our reporter system possibly combined with wild-type challenge constitute the ideal model to investigate the earliest aspects of sexual transmission, potentially advancing the development of next generation PrEP approaches.

## MATERIALS AND METHODS

### Ethics statement

All animal studies were conducted in accordance with protocol P0153, approved by Northwestern University and Tulane National Primate Research Center Institutional Animal Care and Use Committees (IACUC). This study was carried out in accordance with the *Guide for the Care and Use of Laboratory Animals* of the National Institutes of Health and the recommendations of the Weatherall report, *The Use of Nonhuman Primates in Research*. Animals were euthanized by sedation with ketamine hydrochloride injection followed by intravenous barbiturate overdose, in accordance with the recommendations of the panel on Euthanasia of the American Veterinary Medical Association.

### Dual-reporter viral vector—design and production

Our SIV-based pseudovirus dual-reporter vector (Fig. 1A) was generated by modification of the SIV3^+^ vector [40]. The two reporters, codon-optimized firefly luciferase and iRFP670 are separated by IRES; iRFP670 was chosen for its unique narrow emission spectrum (670 nm) [15]. To ensure robust expression, transcription of both reporters was driven by the constitutive immediate-early CMV promoter and stimulated by WPRE [41, 42].

Pseudotyped reporter virus was produced by transfection of 293T cells with four plasmids complexed with polyethyleneimine (Polysciences): dual-reporter vector described above, SIV3^+^ packaging vector, REV expression plasmid DM121, and either a VSVg or JRFL envelope. VSVg pseudotyped particles (VSVg-LI670) were used to validate reporters in culture; JRFL pseudotyped particles (JRFL-LI670) were used in the animal challenge studies. As described previously [2], the 293T cell supernatants containing pseudotyped virus were collected 48 hours post-transfection, purified through 0.22-μm filters, concentrated over 30% sucrose cushions, titrated for infectivity (TCID_50_ range: 10^4^-10^6^) on 293T cells, and stored at −80°C.

### Cell culture

293T cells (American Type Culture Collection) were cultured at 37°C and 5% CO_2_ in Dulbecco’s Modified Eagle Medium (HyClone) containing 10% fetal bovine serum, 100 Um/L penicillin, 100 µg/mL streptomycin and 292 µg/mL l-glutamine (Gibco). Cells were seeded on coverslips in 24-well plates (50,000 cells/well) and, upon reaching 80% confluence, transduced with VSVg-LI670. After four hours, virus-containing supernatant was removed, and cells fed with fresh media. Forty-eight hours later, the transduced cells were fixed, immunostained and analyzed, as described below.

### Non-human primate studies

Four female RMs were used for this study. At the Tulane National Primate Research Center, each animal received four to six rectal biopsies at random, and were then immediately inoculated rectally with 5 mls concentrated JRFL-LI670 virus (TCID_50_ range: 10^4^-10^6^). Animals were sacrificed 48 hours after inoculation (the shortest time interval in which we could reliably detect reporter expression). Immediately, anorectal tissue encompassing the first 6 to 7 cm of the distal colon was extirpated in one piece, and shipped overnight in RPMI medium on ice for further processing in the Hope Laboratory at Northwestern University.

### In vivo imaging system (IVIS)

We employed IVIS to rapidly and efficiently screen the large pieces of anorectal tissue for a relatively small number of transduced cells. The anorectal tissue piece was first washed gently, yet thoroughly, in PBS to remove as much fecal matter as possible and placed in the PerkinElmer IVIS Lumina Series III device to image background luminescence. Disposable tissue biopsy punchers were used to excise the visible biopsy sites (3 mm across), and the remaining tissue was cut into six large pieces. All tissue pieces were incubated in 100 mM d-luciferin (Biosynth) for 10 minutes, and IVIS imaged a second time—in this instance, for luciferase expression. One or more pieces of anorectal tissue with the highest expression (ie, presumably the highest number of transduced cells), were cut into six smaller pieces (2 mm^2^) and re-imaged by IVIS, to even further isolate the areas of the tissue with the highest lucerifase expression. All luminescent signals were analyzed using Living Image Software (PerkinElmer). In this manner, we narrowed down the areas of interest for the follow-up fluorescent microscopy. However, all pieces of tissue were snap frozen in optimal cutting temperature (OCT) compound.

### Immunolabeling and microscopy

OCT blocks of anorectal tissue were cryosectioned (20-µm), fixed in 1% formaldehyde in PIPES buffer (Sigma) and blocked with normal donkey serum (Jackson Immuno). Cryosections and 293T cells were immunostained for luciferase with polyclonal rabbit anti-firefly luciferase antibody (clone ab21176, abcam) pre-labeled with Zenon Alexa Fluor 488 Rabbit IgG Labeling Kit (Thermo Fisher Scientific). In addition, cryosections were immunolabeled with one or more of the following four antibodies: 1) mouse anti-human CD4 (clone OKT4, hybridoma supernatant) with rhodamine Red-X donkey anti-mouse IgG (Jackson Immuno); 2) rabbit anti-human CD3 (clone SP7, abcam) with rhodamine Red-X donkey anti-rabbit IgG (Jackson Immuno), 3) mouse anti-human CD196 prelabeled with Alexa Fluor 488 (CCR6 marker; clone G034E3, BioLegend) and 4) E-Cadherin (BD Pharmagin, catalog number 560062). Secondary-antibody-only negative-control images were used to set specificity thresholds for each channel, when appropriate. After antibody incubations, cryosections (and 293T cells) were incubated with Hoechst 33342 fluorescent stain (Thermo Fisher Scientific) to visualize nuclei. Image stacks (20 to 40 sections in the Z plane in 0.5-µm steps) were acquired and deconvolved using SoftWoRx software on a DeltaVision Elite (GE) inverted microscope equipped with a 60x magnification objective.

Statistical analyses were performed focusing on the infected cells, proportions averaged over the four monkeys were used in a goodness of fit test to examine if at least one of the proportions is different from the others (alpha=0.05). If at least one proportion was different, 6 pairwise goodness of fit tests would be performed to determine which groups are different, using a Bonferroni corrected alpha=0.008.

### Fluorescent in situ hybridization (FISH)

To detect *RORc* transcript variants encoding RORγT, CAL Fluor Red 610-conjugated RNA probes were generated using Stellaris RNA FISH probe designer (BioCat). Cryosections (20 µm) were fixed with 3.7% (vol/vol) formaldehyde in 1x PBS blocked with NDS, and then double immunolabeled with CD3 and CCR6 antibodies, as outlined above (Immunolabeling and microscopy). Sections were then incubated in 70% ethanol for 10 minutes at room temperature, and then FISH performed, according to the manufacturer’s recommendations for sequential IF and FISH (Biosearch Technologies); attention was taken to minimize exposure of tissue and reagents to RNases. Imaging was performed as outlined above (Immunolabeling and microscopy).

### Spectral imaging

Spectral imaging was performed on immunolabeled cryosections using a Nikon A1R laser scanning confocal microscope equipped with four laser lines (408, 488, 561, and 638), a 60x magnification objective, Nikon Elements software, and a 32-channel PMT spectral detector coupled with software-based linear un-mixing algorithm. Imaging area was set in the x-y dimension using a region of interest (ROI) feature. After adjusting intensities and exposure times for the four lasers, the pre-selected ROI was imaged at various intervals along the wavelength axis to establish the pattern of intensity changes at different emission bands. In this way, we could determine the emission spectrum for each fluorophore by plotting pixel intensity vs center wavelength for each emission band. Depending on the experimental design, we imaged up to four fluorophores at a time. The fluorophores and their excitation/emission spectra (in nm) were as follows: Hoechst (345/355), Alexa Fluor 488 (495/519) for luciferase, rhodamine Red-X antibody (560/580) for CD4 and CD3, and direct fluorescence of iRFP670 (643/670).

### Nested PCR

Genomic DNA was isolated from 3 to 5 mg of frozen tissue using the DNeasy Blood & Tissue Kit (Qaigen). To detect a 244-bp DNA fragment of the luciferase reporter gene, each nested PCR reaction used 250 ng of genomic DNA, a published amplification procedure,[2] and an iCycler Thermal Cycler system (Bio-Rad). First-round primers were 5′-GAAGCGCTATGGGCTGAATA-3′ (forward) and 5′-GTCGTACTTGTCGATGAGAGTG-3′ (reverse). In the second round, 2 µl of first-round reaction product was amplified with 5′-CATGGATAGCAAGACCGACTAC-3′ (forward) and 5′-GATGATCTGGTTGCCGAAGA-3′ (reverse). Each DNA sample was tested in 24 or more replicates. Negative-control unchallenged rectal tissue was from animals not exposed to the luciferase-containing reporter vector. Second-round PCR products were separated in 1.5% agarose gels and visualized by ethidium bromide staining. For each lane, sequences were confirmed by extracting DNA with the QIAquick Gel Extraction Kit (Qaigen) with the second-round primers.

## ACKNOWLEDGEMENTS

We wish to thank Edgar Matias for help with tissue sectioning, Edward Allen for virus production, Gianguido C. Cianci and Elena Martinelli for valuable discussions and for reading the manuscript, and Andrea Cimarelli for kindly providing us with SIV3^+^ vector.

Authors have no conflict of interest to declare.

DM designed and performed the experiments, prepared the figures and wrote the manuscript. WAG generated LI670 construct and performed the nested PCR analysis. NG aided in the IF and RNA-FISH analysis. MDM coordinated animal challenges, virus collection and helped with tissue processing. AJF performed the statistical analyses. RSV conducted animal challenges and supervised animal necropsies. TJH designed the study and provided feedback at all stages. All of the authors were involved in reading and editing the manuscript and consented to its submission.

## FIGURE LEGENDS

**FIG S1.**
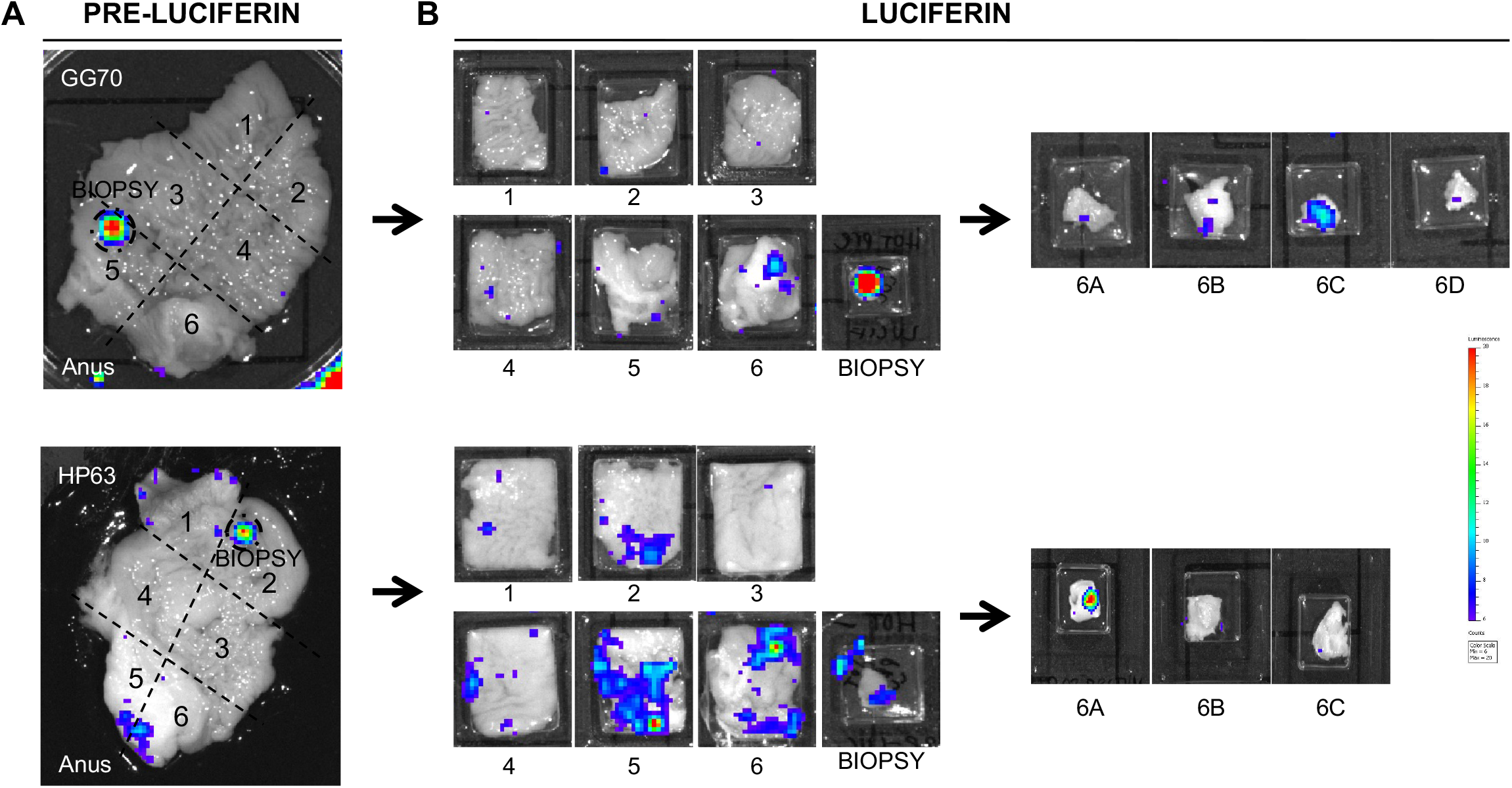
Luciferase reporter expression in anal and rectal tissue of vector-inoculated RMs. Animals were inoculated with JRFL-pseudotyped dual-reporter vector, sacrificed 48 hours later and then a large area of anorectal tissue was removed in one piece. (A) Extirpated anorectal tissue from two animals (GG70 and HP63) imaged by IVIS to assess background luminescence; biopsy sites and anal pole are indicated. (B) Tissue at the biopsy sites was excised and remaining tissue cut into six large pieces as indicated by dotted lines in A; all pieces were soaked in d-luciferin and imaged by IVIS to visualize luciferase expression. For each animal, the tissue piece with the highest expression was cut into smaller pieces that were then re-imaged by IVIS.

**FIG S2.**
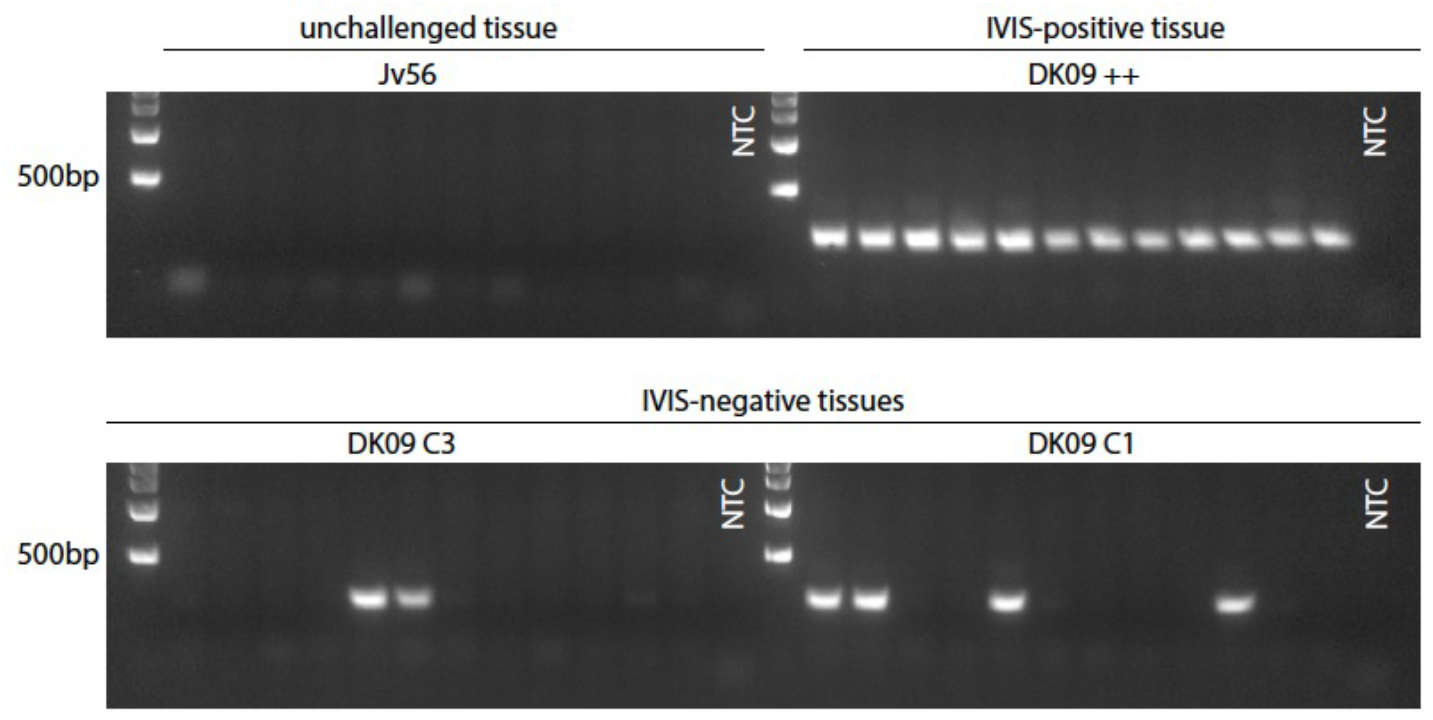
Nested PCR validates IVIS-luciferase data, while also revealing additional transduction events. To detect a 244-bp fragment of the firefly luciferase reporter gene, genomic DNA (template) was extracted from cryosections of anorectal tissue from RM DK09, including positive-control DNA from biopsy site tissue with robust luciferase expression (Figs. 2, S1). Negative-control DNA was extracted from tissue from an unchallenged, naïve RM from an unrelated study. Each lane represents a unique DNA sample from a unique cryosection.

**FIG S3.**
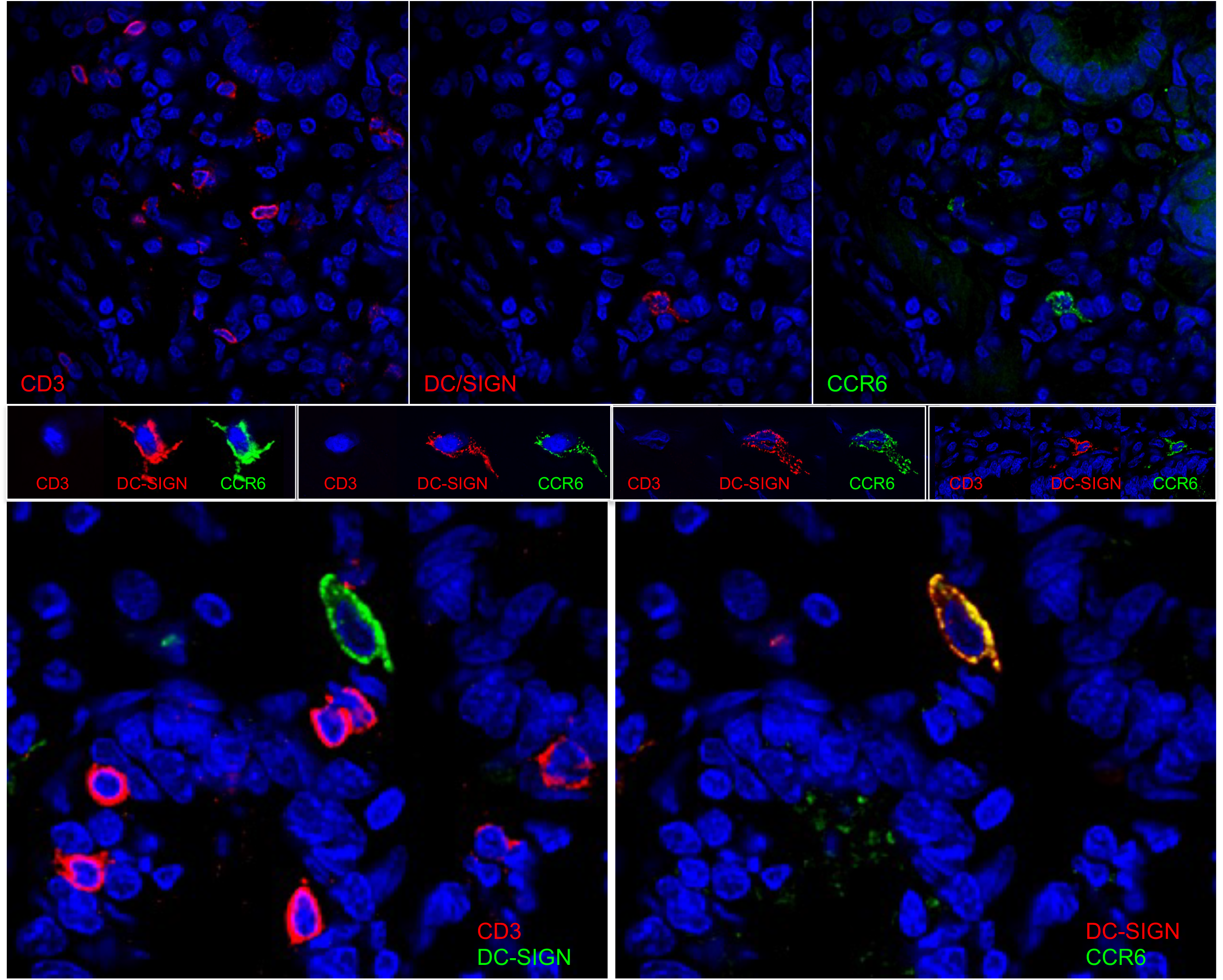
All of CCR6^+^ CD3^−^ cells express DC/SIGN molecule CD209 in the anorectal tissue. Cryosections were immunolabeled with CD3, CD209 (DC/SIGN) and CCR6 antibodies and Hoechst stain. Each row shows images of the same field. The top panel shows a low magnification (40x) to show distribution of T cells and iDCs throughout the tissue, while middle and low bottom images show close up magnification to show distinct morphology of iDCs (larger size with multiple dendrite projections) vs. T cells (small granular in appearance).

**FIG S4.**
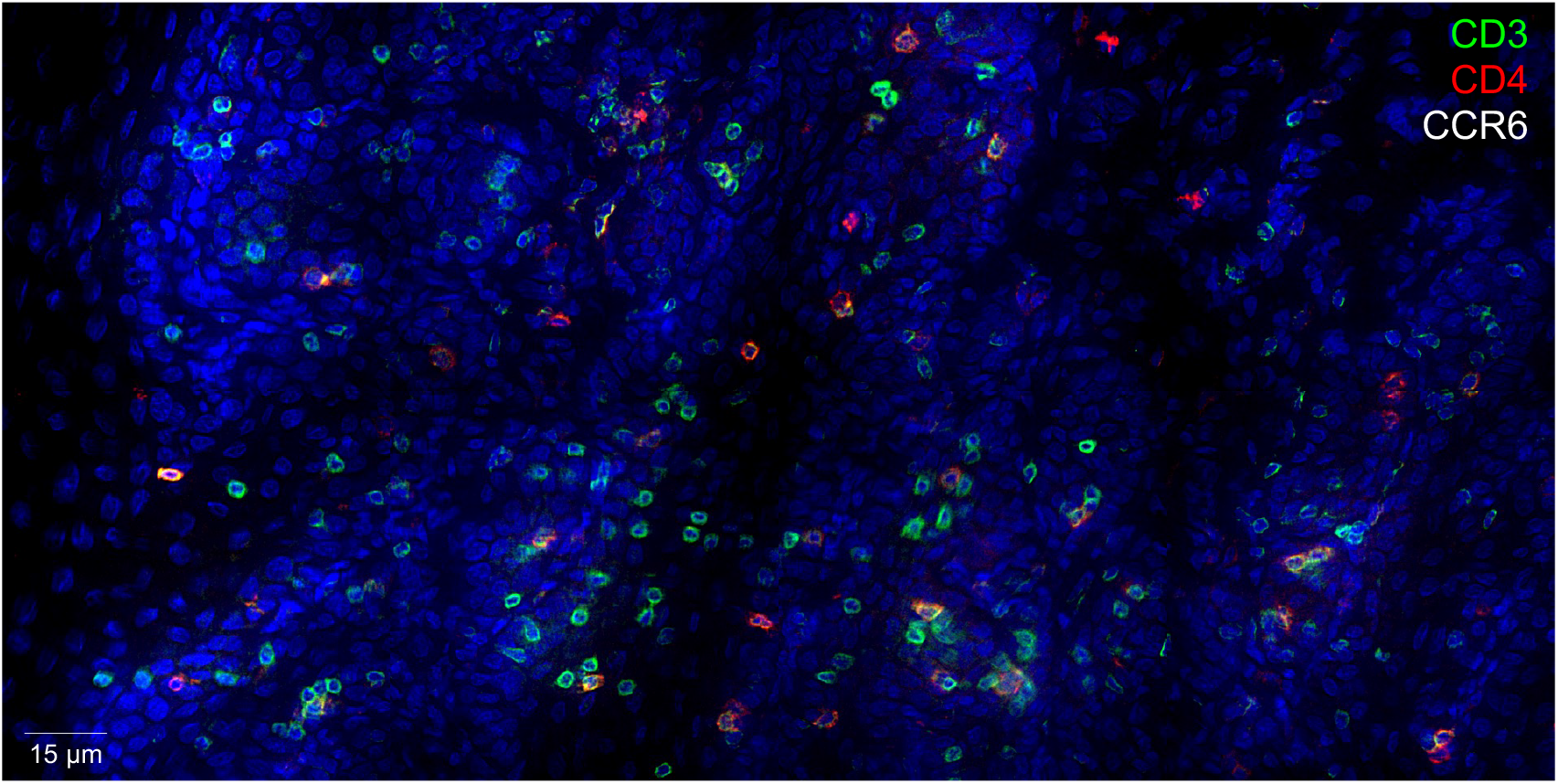
Relative abundance and distribution of CD3^+^, CD4^+^ and CCR6^+^ cells in anorectal tissue. Montage of multiple low magnification (40x) images of an immunolabeled and Hoechst-stained cryosection depicting a large swath of anorectal tissue.

---

**Table S1.**
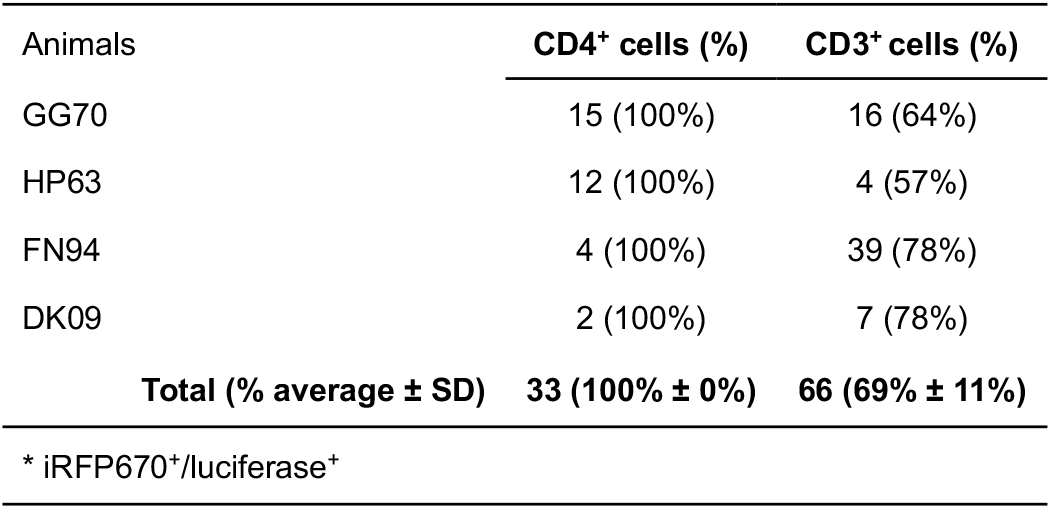
CD4 and CD3 phenotypic analysis of transduced* cells after anorectal inoculation with JRFL-pseudotyped virions.

| Animals | CD4<sup>+</sup> cells (%) | CD3<sup>+</sup> cells (%) |
| --- | --- | --- |
| GG70 | 15 (100%) | 16 (64%) |
| HP63 | 12 (100%) | 4 (57%) |
| FN94 | 4 (100%) | 39 (78%) |
| DK09 | 2 (100%) | 7 (78%) |
| **Total (% average ± SD)** | **33 (100% ± 0%)** | **66 (69% ± 11%)** |
| * iRFP670<sup>+</sup>/luciferase<sup>+</sup> |  |  |

**Table S2.** Frequency analysis of four subtypes of CD4^+^ cells in anorectal tissue.

| Animals | Cell Count (%) |  |  |  |  |
| --- | --- | --- | --- | --- | --- |
|  | CD3<sup>+</sup> |  | CD3<sup>-</sup> |  | TOTAL |
|  | CCR6<sup>+</sup> | CCR6<sup>-</sup> | CCR6<sup>+</sup> | CCR6<sup>-</sup> |  |
| GG70 | 9 (6%) | 110 (70%) | 5 (3%) | 32 (20%) | 156 |
| HP63 | 15 (9%) | 116 (73%) | 2 (1%) | 26 (16%) | 159 |
| FN94 | 21 (10%) | 172 (79%) | 2 (1%) | 22 (10%) | 217 |
| DK09 | 20 (18%) | 67 (60%) | 3 (3%) | 21 (19%) | 111 |
| **Total (% average ± SD)** | **65 (12% ± 5%)** | **465 (71% ± 9%)** | **12 (2% ± 1%)** | **101 (15% ± 4%)** | **643** |

